# The evolution of hematopoietic cells under cancer therapy

**DOI:** 10.1101/2020.10.29.360230

**Authors:** Oriol Pich, Albert Cortes-Bullich, Ferran Muiños, Marta Pratcorona, Abel Gonzalez-Perez, Nuria Lopez-Bigas

## Abstract

Chemotherapies may influence the evolution of somatic tissues through the introduction of genetic variation in cells and by changing the selective pressures they face. However, the contributions of chemotherapeutic agents to the mutation burden of healthy cells and to clonal expansions in somatic tissues are not clear. Here, we exploit the mutational footprint of some chemotherapies to explore their influence on the evolution of hematopoietic cells. Cells of Acute Myeloid Leukemia (AML) secondary to treatment with platinum-based drugs showed a clear mutational footprint of these drugs, indicating that healthy blood cells received chemotherapy mutations. In contrast, no trace of 5-fluorouracil (5-FU) mutational signature was found in AML secondary to exposure to 5-FU, suggesting that cells establishing the AML were quiescent during treatment. We used the platinum-based mutational signature as a barcode to precisely time clonal expansions with respect to the moment of exposure to the drug. The enrichment for clonal mutations among treatment-related mutations in all platinum-treated AMLs shows that these secondary neoplasms begin their clonal expansion after the start of the cytotoxic treatment. In contrast, the absence of detectable platinum-related mutations in healthy blood samples with clonal hematopoiesis is consistent with a clonal expansion that predates the exposure to the cytotoxic agent, which favours particular pre-existing clones.

## Introduction

Somatic tissues evolve as a result of the interplay between genetic variation ---contributed by a range of endogenous and external mutational processes--- and selective constraints acting at the level of organs or tissues^1–3^. Chemotherapies cause the death of large amounts of cells, thus imposing specific selective constraints on somatic tissues^4^. Certain cells, able to withstand chemotherapies by virtue of certain advantageous mutations or phenotypic characteristics may subsequently expand to replenish the exhausted tissue after the insult is withdrawn. Some widely used chemotherapies, due to their mutagenic mechanism, also contribute to the genetic variation present in exposed tumor cells or cell lines^5–10^. Nevertheless, it is not known whether these chemotherapeutic agents leave their mutational footprint in healthy cells. Furthermore, the immediate effects of the exposure to chemotherapies ---whether they leave a footprint or not--- on the evolution of healthy tissues is not clear.

One homeostatic process in which the long-term effects of chemotherapies have been extensively studied is hematopoiesis. It is known, for example, that secondary hematopoietic malignancies (such as AML) occur in patients who are exposed to chemotherapies as part of the treatment of a solid malignancy^11,12^. Moreover, clonal hematopoiesis (CH), a condition related to aging across the human population is known to occur more frequently among people previously exposed to chemotherapies^13–19^. CH, in turn, is associated with other health risks, such as hematopoietic malignancies or increased incidence of cardiovascular disease^13,17,20–22^. The molecular mechanisms underlying the advantage provided by some CH-causing mutations affecting DNA-damage and repair genes in the face of certain chemotherapies have been unraveled^16^. However, the precise timing of the clonal expansion that ultimately causes treatment-related CH or treatment-related AML with respect to the exposure to chemotherapy remains elusive. This is key to understanding whether the cytotoxic agent may be the cause of this clonal expansion or only provides a new evolutionary constraint that favors a pre-existing clone.

Chemotherapy-related mutations are only detectable through bulk sequencing when one or several of the cells where they have occurred expand. Thus, we reasoned that their detection in therapy-related AML (tAML) and CH cases provides a powerful tool to study the evolution of such conditions. Specifically, we set out to determine whether cells that were healthy at the time of their exposure to chemotherapeutic agents bear their mutational footprint. Furthermore, we used this footprint as a barcode^23^ to precisely time the clonal expansion with respect to the moment of exposure to the drug. Ultimately, this allowed us to study how chemotherapies interfere with healthy hematopoiesis, causing the emergence of treatment-related AML (tAML) or treatment-related CH (tCH).

## Results

### The role of chemotherapies in the evolution of tAMLs

We and others previously observed that certain chemotherapies, through direct DNA damage or interference with the replication machinery leave a mutational footprint in the metastatic tumors of patients exposed to them as part of the cancer treatment^5,7^. We reasoned that such chemotherapies, being systemic, may also be able to leave their mutational footprint in healthy cells. However, detecting private chemotherapy mutations ---in the absence of any clonal expansion--- in healthy cells is extremely challenging. We reasoned that secondary neoplasms could give us the opportunity to study this. Secondary neoplasms, such as tAMLs, which appear in some patients following treatment of a primary solid tumor, originate from hematopoietic cells that were healthy at the time of exposure (Fig. 1a). It is not clear whether these tAMLs are driven specifically by drug-related mutations, or if they appear as a consequence of the evolutionary bottleneck chemotherapies posed on hematopoiesis, or as a contribution of both factors.

**Figure 1.**
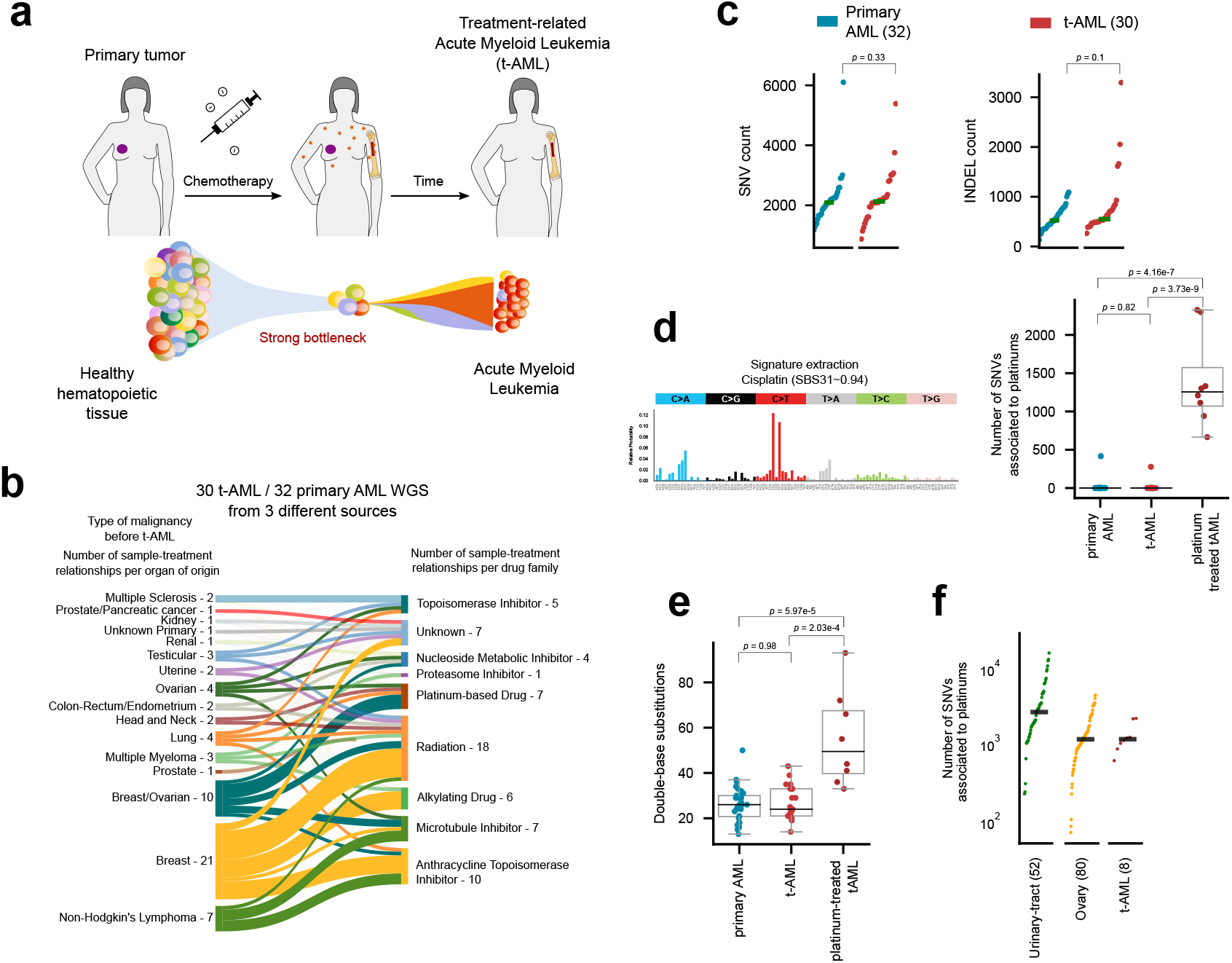
The mutational footprint of chemotherapies across treatment-related AMLs. a) Healthy hematopoietic cells at the time of exposure to a chemotherapy are faced with a bottleneck that reduces the population, leading to the development of AML over time. b) The whole-genome sequence of thirty tAML cases of patients who suffered from a primary solid tumor and were treated with different anticancer drugs (represented in the sankey diagram) were obtained from three different sources, including three cases sequenced in-house. These were analyzed in combination with 32 cases of primary AMLs (WGS AML cohort). c) Burden of single nucleotide variants and indels does not statistically differ across cases of the WGS AML cohort (two-tailed Mann-Whitney *p*=0.33, and 0.1 respectively). d) Mutational profile of a platinum-related signature active across cases in the WGS AML cohort. All tAML cases from patients exposed to platinum-related drugs exhibit activity of the signature (two-tailed Mann-Whitney *p*=3.37e-9, and 4.16e-7). e) The number of double base substitutions of platinum-exposed tAML cases is significantly higher (two-tailed Mann-Whitney *p*=5.97e-5 and 2.03e-4) than that of primary AMLs or tAMLs from patients exposed to other anticancer therapies. f) Number of mutations contributed by the platinum-related signature across tAML cases in comparison with that counted across metastatic tumors from several organs of origin.

To explore the role of anti-cancer treatments in the development of tAMLs, we collected a cohort of 30 (3 in-house) whole-genome sequenced tAMLs (Fig. 1b; Fig. S1a) and 32 primary AMLs (WGS AML cohort)^24,25^. Overall, no significant differences are appreciable in total somatic SNVs (*p*=0.33) or indels (*p*=0.1) burden between tAMLs and primary AMLs, as previously reported^24^ (Fig. 1c; Table S1). However, all tAMLs from patients exposed to platinum-based chemotherapies exhibit a mutational footprint associated with these drugs (Fig. 1d; Fig. S1b; Table S1). This signature is highly similar (cosine similarity 0.94) to a platinum-related signature identified across metastatic tumors from patients exposed to the drug^5,26^. (Heretofore, we refer to this mutational signature either as platinum-related.) Moreover, they harbor a significantly higher number of double base substitutions than tAMLs of patients exposed to other drugs and primary AMLs, as expected from the mutagenic mechanism of platinum-based drugs^27^ (Fig. 1e; Table S1). Despite the profound phenotypic differences that exist between healthy hematopoietic cells and solid tumor cells, the number of platinum related mutations in tAMLs is similar to that detected across metastatic tumors from patients exposed to the same chemotherapies^5^ (Fig. 1f). Therefore, the exposure to platinum-based drugs appears to be a sufficient condition for the acquisition of platinum-related mutations in healthy blood cells.

Interestingly, no trace of the 5-FU-related mutational footprint appears in any of the three tAML cases from patients exposed to the drug (Fig. S1b). Platinum-based drugs directly damage the DNA of cells generating bulky adducts. Conversely, 5-FU through the inhibition of thymidylate synthase^28^, alters the pool of nucleotides available for DNA synthesis, and as analogous to thymidine, could be incorporated to nascent DNA strand. We hypothesized that cells that divide during the treatment will incorporate 5-FU mutations, whereas cells that remain quiescent during the exposure to 5-FU will not. The lack of the 5-FU mutational footprint would indicate that cells that establish tAMLs were quiescent (potentially hematopoietic progenitors) at the time of treatment^29–31^.

In summary, we demonstrated that platinum-based therapies leave their characteristic mutational footprint across healthy hematopoietic cells upon exposure to them. Intriguingly, the footprint that characterized 5-FU appears to be absent from these exposed healthy hematopoietic cells, possibly because quiescence is a key mechanism of survival.

### Footprint of chemotherapies with differing mutagenic mechanisms

To pursue the hypothesis that the differing mutagenic mechanism of platinum-based drugs and 5-FU underlies the differences in detection of their mutational footprint, we resorted to a cohort of metastatic tumors^32^ (metastasis cohort) among which we and others had previously identified the mutational footprint of both drug families^5,7^ (Fig. S2a). As part of their treatment the patients in this cohort were exposed to chemotherapies for a period of time following the detection of their primary tumors (**time of treatment**) and some time after the completion of these treatment regimens metastatic tumors were biopsied (**time since the end of treatment**). In order to be able to detect mutations of platinum-based drugs or 5-FU in metastatic tumors two conditions are necessary. First, cells exposed to the drugs must acquire treatment-related mutations; second, some of these cells with treatment mutations must subsequently expand to a level that is sufficient for the mutations to rise above the limit of detection of bulk sequencing (Fig. 2a). Thus, treatment mutational signatures could be used to study the state of cells at the time of exposure to the chemotherapy and the level of clonal expansion after treatment. Indeed, an important fraction of metastatic tumors from patients exposed to platinum-based drugs or 5-FU did not exhibit the mutations related to these drugs (Fig. 2b). As cancer treatment is a relatively recent event in the life of a patient, chemotherapy-induced mutations tend to be more subclonal than those contributed by other mutational processes^5^. However, the fraction of treatment mutations that are clonal across metastatic tumors varies widely between tumors (Fig. S2b), with a subset of metastases in which more than half of all treatment mutations are clonal. The fraction of cells in the tumor sharing chemotherapy-induced mutations can thus be used as a measure of the strength of the evolutionary bottleneck and clonal expansion posterior to treatment. For instance, the enrichment for chemotherapy-induced clonal mutations in a subset of cases, is indicative that the metastatic clonal expansion has taken place between the start of the treatment and the biopsy of the metastasis. This is compatible with, for example, a metastasis seeding posterior to the acquisition of treatment mutations.

**Figure 2.**
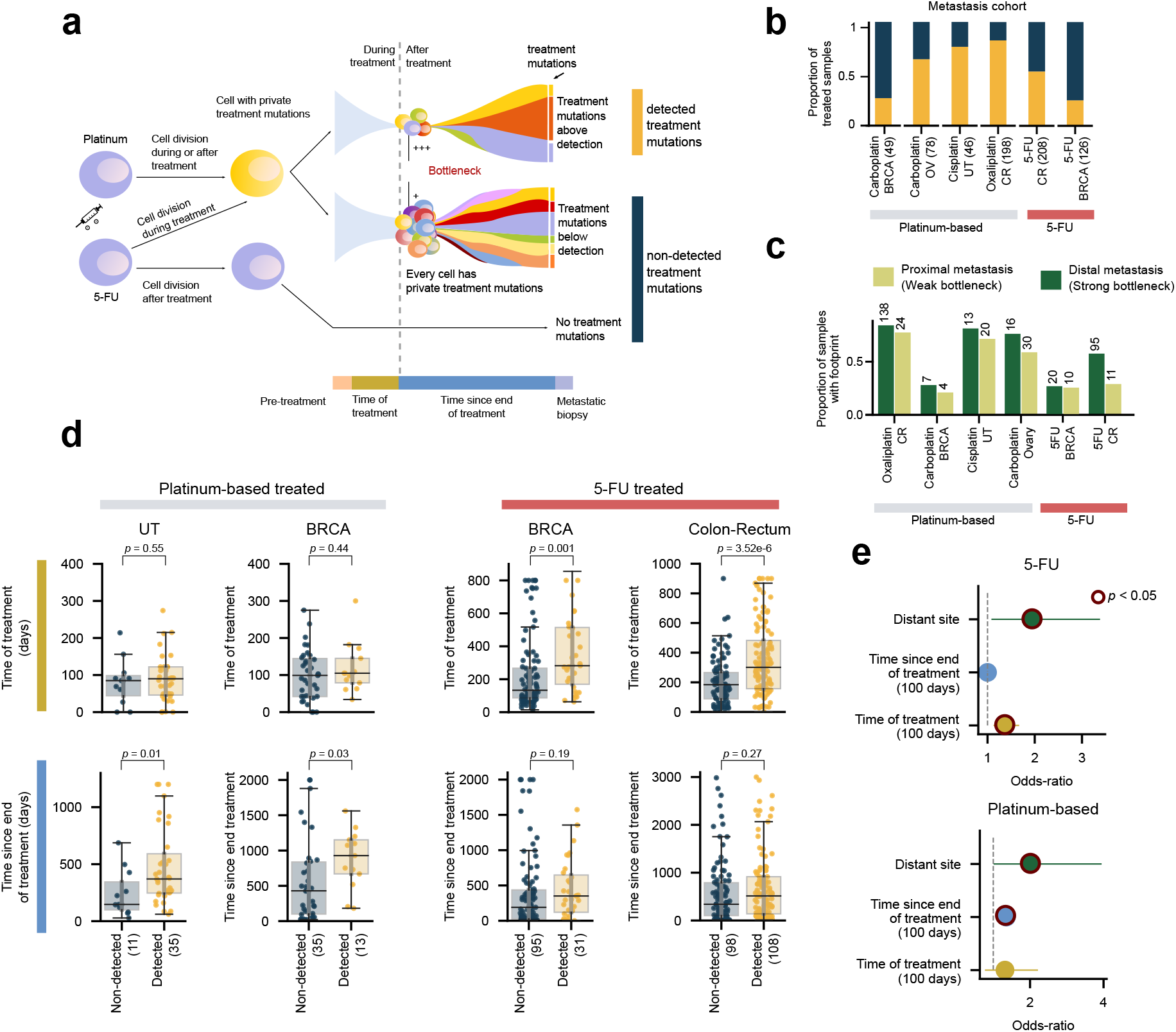
Different mutagenic mechanisms of platinum-based drugs and 5-FU. a) Platinum-based drugs and 5-FU contribute mutations in the DNA via two different mechanisms (illustrated by the blue cells in the left panel). While the former creates adducts on the DNA, the latter alters the pool of nucleotides available for DNA replication. Thus, cells exposed to platinum-based drugs will carry mutations of the treatment after DNA replication, irrespective of whether cells are quiescent at the time of exposure or not (top arrow). On the other hand, 5-FU-exposed cells will only incorporate mutations if their DNA is replicated while the pool of nucleotides is still distorted by the drug (middle arrow). If it has been restored before DNA replication, no mutations will be incorporated (bottom arrow). Therefore, immediately following exposure, two scenarios are possible: the surviving cells either bear treatment mutations (yellow cell) or not (blue cell). We reasoned that whether the treatment mutations are visible at the time of metastasis depends on the strength of the evolutionary bottleneck facing the primary tumor and its timing with respect to the exposure (see periods of time represented in different colors below figure). A stronger bottleneck during or immediately after treatment will lead to treatment mutations detectable through bulk sequencing (top drawing). On the contrary, a weaker bottleneck, or a bottleneck occurred before treatment (for example, if the metastasis predates the treatment) will lead to treatment mutations below the limit of detection of the bulk sequencing (middle drawing). b) Proportion of samples from metastatic tumors with different organs of origin taken from patients exposed to platinum-based drugs or 5-FU with detectable treatment mutations. c) Proportion of samples with detectable treatment mutations among distant or proximal metastases. d) Distribution of time of treatment (days) and time after treatment (days) of samples of metastatic tumors of different organs of origin with detectable or undetectable treatment mutations taken from patients exposed to platinum-based drugs or 5-FU. Groups of metastatic tumors were compared using the one-tailed Mann-Whitney test. e) Results of logistic regressions showing the influence of different variables on the detection of platinum-based drugs (bottom) or 5-FU (top) related mutations across metastatic tumors. Variables that significantly influenced the likelihood of detection are circled.

We also observed that the mutational footprint of both platinum-based drugs and 5-FU is detected more often in distant metastases than in proximity metastases (Fig. 2c,e). Distant metastases are known to undergo a stronger evolutionary bottleneck^33^. Therefore, it is more likely in distant metastases that treatment mutations appear above the limit of detection of the bulk sequencing. In addition, significant differences in the time since the end of treatment distinguished tumors with the footprint of platinum-based drugs from those with no footprint (Fig. 2d,e; Fig. S2c; Table S2). The longer the time between the end of the treatment and the biopsy of the metastasis the more likely the cells seeding it has experienced a full clonal expansion, and thus the higher the odds to detect the footprint.

Intriguingly, tumors with detectable footprint of 5-FU were exposed to the drug significantly longer than those without the footprint, but both groups showed no significant differences in the time since the end of treatment (Fig. 2d,e; Fig. S2c; Table S2). Unlike unrepaired DNA adducts left by platinum-based drugs, which may be directly converted into mutations as DNA replicates, 5-FU-generated mutations can only appear if, at the time of replication, the nucleotide pool has not been restored. This is more likely to occur the longer cells are exposed to 5-FU, and is probably the reason why the time of treatment is key to observing 5-FU mutations in the metastases. This would also explain why there are no significant differences in the time after the treatment for the metastases that show the 5-FU footprint and those that do not. The significantly longer time of treatment of tumors with the 5-FU footprint may account for the time required for the full clonal expansion.

In summary, the level of clonality of treatment-related mutations is variable across metastasis. Treatment-related mutations are only detected in a fraction of the tumors in the metastasis cohort. In these tumors, the mutational footprint of the chemotherapy may be used as a barcode^23^. The detection of these footprints provides clues about the state of cells during treatment. It also implies that the treatment preceded the beginning of the clonal expansion giving rise to the metastasis. Therefore, as stated above, the absence of a detectable 5-FU mutational footprint in tAMLs may be due to the malignancy being started by hematopoietic cells that were quiescent during their exposure to the drug.

### The effect of chemotherapies on the selective constraints faced by hematopoietic cells

Next, we used the knowledge leveraged in metastatic tumors to ask whether the clonal expansion of the tAML is prior or posterior to the exposure of patients to platinum-based drugs. We found that, as expected, across the WGS AML cohort, platinum-related mutations are more active among clonal mutations than those contributed by other mutational processes (Fig. 3a). This is consistent with the treatment predating the clonal expansion of the hematopoietic cells that founded the tAML.

Across both, tAMLs and primary AML cases, the most prevalent mutational signature is associated with the steady accumulation of mutations in hematopoiesis (HSC signature^34^) (Fig. 3b,c; Fig. S3a; *p*=2.86e-17). The linear relationship between the number of mutations contributed by this process of hematopoiesis and age, which has been observed in healthy hematopoietic cells is maintained across primary AMLs, albeit with a slight acceleration (Fig. 3c; *p*=2.34e-11). The mutations contributed by other processes active in hematopoietic cells do not accumulate linearly with age (Fig. S3b). However, the linear relationship between the gain of hematopoiesis-related mutations and age is lost in tAMLs, and apparently, this is independent of whether or not the chemotherapy leaves a mutational footprint (Fig. 3d, *p*=0.81). This suggests that regardless of their mutagenic effect, chemotherapies alter the developmental dynamics of the hematopoietic compartment^30,35^.

**Figure 3.**
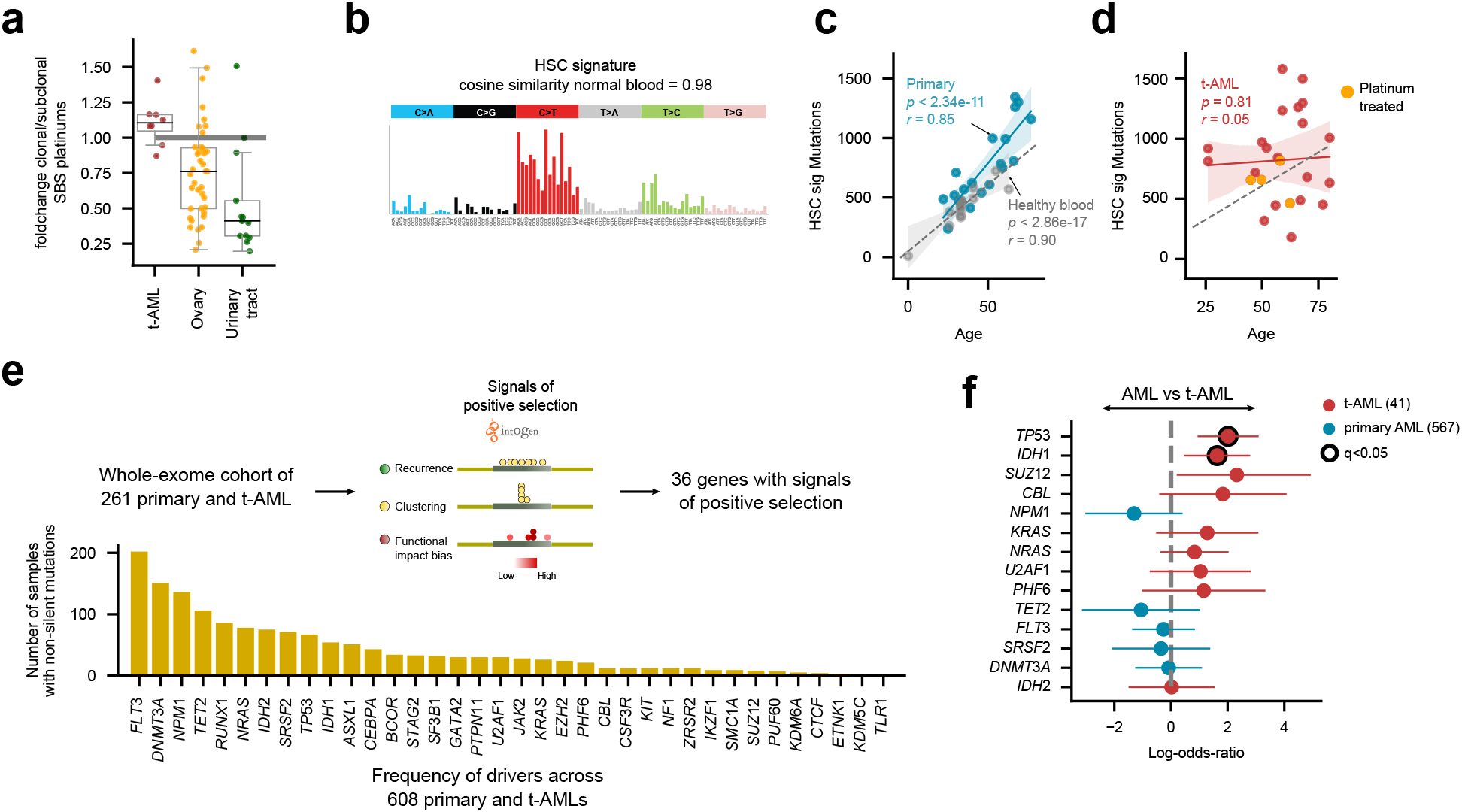
The development of treatment-related AMLs. a) Relative activity of platinum related mutations among clonal mutations with respect to subclonal mutations across tAMLs and the metastatic tumors of several organs of origin. b) Mutational profile of the HSC signature active across tAML cases. c) Linear relationship between the age of patients and the number of HSC mutations of primary AMLs (blue dots). The gray dots correspond to healthy blood donors and thus the regression of their HSC mutation burden to the age represents the normal accumulation of mutations in the process of hematopoiesis. In the figure *r* represents the Pearson#8217;s correlation coefficient and *p*, its associated p-value. d) No significant regression is obtained between the HSC mutation burden and the age of patients with tAML. e) Genes with detectable signals of positive selection across the subset of 261 AMLs of the WES AML cohort with a healthy control sample. The recurrence of mutation shown by the bars is measured across the entire cohort. f) Overrepresentation of mutations of different genes across primary or treatment-related AMLs. Significant cases (Benjamini-Hochberg FDR<0.05) appear circled.

Since the hematopoietic cells founding the primary and tAMLs face different selective constraints, we next asked whether mutations in different genes drive both malignancies. To this end, we identified genes under positive selection across 261 whole-exome sequenced AMLs, 16 of them secondary to treatment. These are part of a cohort of 608 AML patients^36^, 41 of which correspond to tAML cases (WES AML cohort; Fig. S3c). We identified signals of positive selection in the mutational patterns of 36 genes across this cohort (Fig. 3e; Table S3). Although the same set of genes drive both primary and tAML cases, some differences are appreciated in the prevalence of their mutations (Fig. S3d; Table S3), such as an enrichment of TP53 mutations among tAMLs ---an effect previously observed in a smaller cohort^24^ (Fig. 3f). Interestingly, we observed the same for IDH1 mutations. On the contrary, mutations of NPM1 are underrepresented in cases of tAML.

### The evolution of treatment-related clonal hematopoiesis and tAML

Clonal hematopoiesis (CH) is a condition characterized by the presence in the blood of individuals of an expanded HSC clone, driven by advantageous somatic mutations^13–17,19,37,38^. In the growth conditions of the healthy bone marrow niche, or faced with particular challenges---such as exposure to cytotoxic therapies--- some of these mutations provide an HSC with a growth advantage with respect to its neighbors. We previously detected the somatic mutations across the blood samples of the ~4,000 donors of the metastasis cohort using the tumor sample as a reference of the germline genome of the donor^38^ (Fig. 4a). The detection of these mutations constitutes evidence of the occurrence of CH in the donors of this cohort. Tracing the signals of positive selection on the mutations observed in genes across donors we have previously obtained a compendium of clonal hematopoiesis (CH) driver genes^38^.

**Figure 4.**
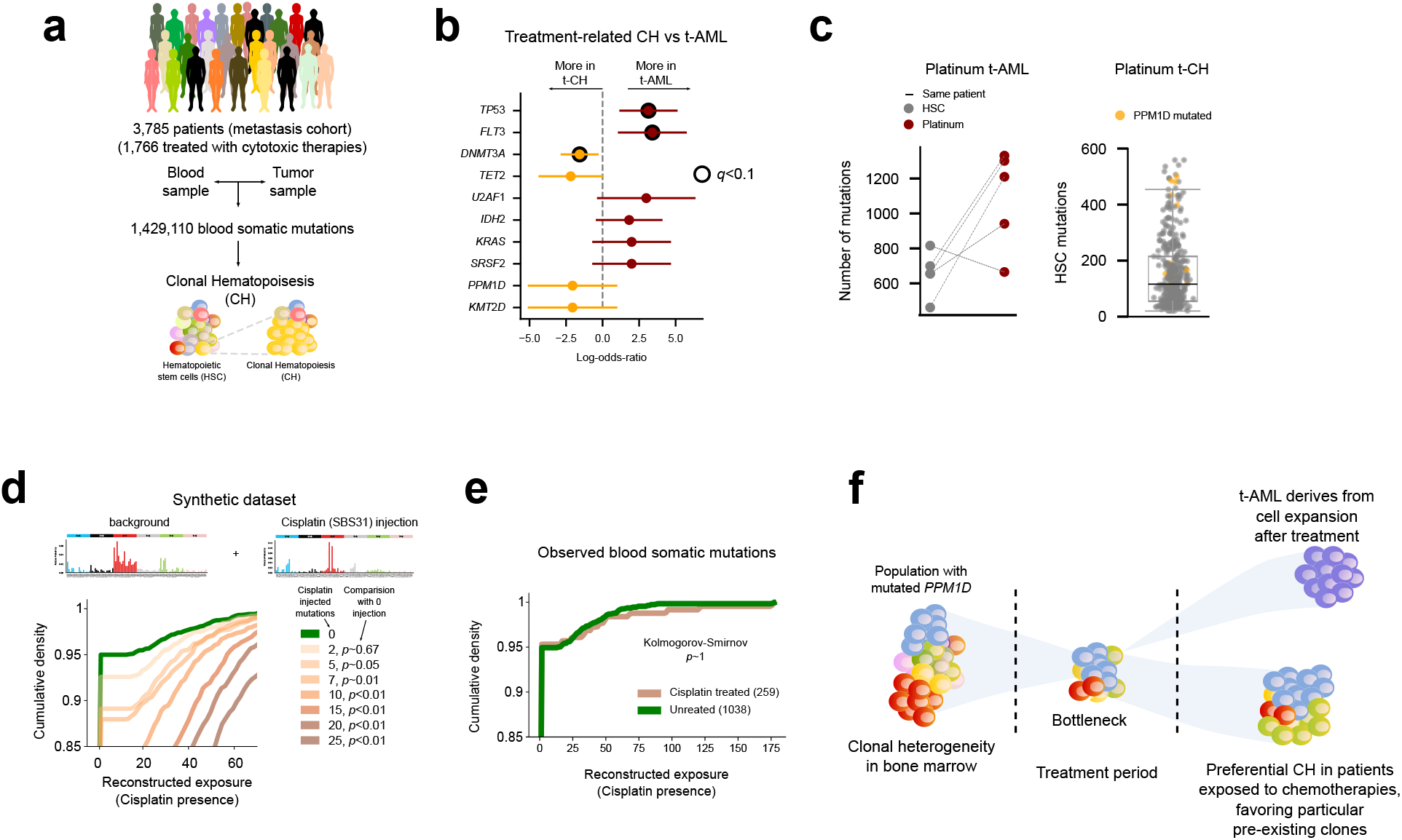
The development of clonal hematopoiesis in chemotherapy-exposed donors. a) Somatic mutations across blood samples from 3,785 patients in the metastasis cohort were detected using the tumor sample as reference of their germline genome. This identification is possible because a process of clonal hematopoiesis has occurred in these samples, rendering the mutations in the founder HSC detectable through bulk sequencing. b) Overrepresentation of mutations in different genes across CH or tAML cases. c) Left panel: number of mutations contributed by the HSC and the platinum-related signature in tAML cases of the WGS AML cohort. Each patient is represented by a straight line joining two circles that represent the contribution of the HSC (gray) and the platinum-related (red) signatures. Right panel: distribution of the number of HSC mutations identified in blood samples across donors in the metastasis cohort. Patients carrying PPM1D mutations are highlighted in yellow. d) To study the detectability of platinum mutations in blood samples of donors of the metastasis cohort, we generated synthetic samples with the same mutational profile. Then, we injected sets of mutations of different sizes that followed the tri-nucleotide probabilities of the platinum-related signature. We then determined the distribution of the reconstructed exposure to this signature of the injected samples. The cohorts of synthetic samples injected with 7 or fewer mutations exhibit a distribution of reconstructed exposure that does not differ significantly from that of the original cohort of synthetic samples. However, the distribution of reconstructed exposure of cohorts of samples injected with 10 or more platinum-related mutations are significantly different from that of the original cohort of synthetic samples. e) No significant differences (Kolmogorov-Smirnov test) are observed in the distribution of the reconstructed exposure (activity) to the platinum-related signature among platinum exposed and platinum non-exposed donors in the metastasis cohort. f) Schematic diagram depicting the possible development of CH and tAML upon chemotherapy exposure of hematopoietic cells.

Exploiting this compendium of CH drivers, we next asked whether mutations in the same genes provide an advantage to hematopoietic cells faced with chemotherapies to develop CH or full-blown tAML. While mutations of TP53, FLT3, IDH1 and other genes driving CH and tAML are overrepresented across tAML cases, DNMT3A mutations appear overrepresented across CH cases (Fig. 4b). It is thus possible that mutations in at least some genes of chemotherapy-related CH do not provide the advantage required to undergo the full clonal expansion that characterizes tAML.

Finally, we reasoned that if, as in the case of tAMLs, the treatment predated the start of the clonal expansion, platinum-related mutations should be detectable in blood samples of donors exposed to these drugs with traces of CH. However, no platinum mutational footprint (or of any drug) is detected through a mutational signatures extraction approach across the blood samples in the metastasis cohort (Fig. S4). To rule out power problems due to the low number of mutations, we then employed mSigAct, a method aimed at recognizing the activity of a particular mutational signature in a dataset of mutations^39^. To first make an estimate of the number of platinum-related mutations expected across blood samples from platinum-exposed donors in the metastasis cohort, we looked at the relationship between platinum-related and HSC mutations across tAMLs. The number of platinum-related mutations across these cases is in the same order of magnitude as the HSC mutations (Fig. 4c, left). Therefore, we predicted that we expect more than 10 platinum-related mutations in the average blood sample from platinum-exposed donors in the metastasis cohort (Fig. 4c, right). Of note, some of the CH cases observed in the cohort have unequivocally expanded against the background of the platinum-based drugs, as evidenced by their PPM1D mutations (yellow-colored in Fig. 4c, right), known to drive the clonal expansion in the presence of these cytotoxic agents^15–17,38^.

Next, we tested the limit of detection of the method and determined that it is able to identify as few as 10 injected platinum-related mutations per sample across a population of synthetic samples with a similar baseline mutational profile and burden as the blood samples of the metastasis cohort (Fig. 4d; Supp. Note). Nevertheless, the application of this method to the blood somatic mutations of the metastasis cohort yielded no trace of the platinum mutational footprint (Fig. 4e; Supp. Note).

The failure to observe the mutational footprint of platinum-based drugs in CH blood samples may thus be explained if the exposure to them does not predate the clonal expansion. We propose that in the majoritary scenario of CH, a variety of clones of hematopoietic cells exists prior to the treatment. Out of these, the clone that carries an advantageous mutation in the bottleneck created by the chemotherapy (such as the PPM1D mutation in patients exposed to platinum-based drugs) is then favored against the background of the selective constraint posed by the chemotherapy, and continues to expand. Conversely, in the case of tAML, with a stronger bottleneck during or after treatment, a clone derived from a single cell gives rise to a leukemic clone (Fig. 4f).

## Discussion

Sequencing somatic tissues provides a window into their evolution through the interplay of genetic variation and selection. The sources of genetic variation and their change throughout time may be resolved by delineating the mutational processes and clonal sweeps to which a tissue has been exposed^26,40^. The detection of the clonal sweeps punctuating the history of a tissue informs on the mutations that provide selective advantages to the constraints faced by its cells^41,42^. The link between the evolutionary dynamics of somatic tissues and the emergence of neoplasias has been known for a long time, and as a result, the interplay of variation and selection has been intensively studied in the context of tumorigenesis^1–3^. Nevertheless, studying the evolution of somatic tissues through life is also key to understanding aging, neurodegenerative diseases and certain cardiovascular conditions. In this endeavour, the resources, approaches and methods developed in recent decades to study cancer emergence and evolution may be repurposed to examine other conditions and diseases.

In this article, building upon our previous finding of the mutational footprint of a group of chemotherapies^5^, we revealed that mutations contributed by platinum-based drugs appear in cells that were healthy at the time of their exposure. Furthermore, the number of treatment mutations that appear in these cells is in the same order of magnitude as those observed in malignant cells after exposure. This exposure appears as determinant, regardless of the phenotypic state of the exposed cell, to observe mutations related to treatments that directly damage the DNA, such as platinum-based drugs^27^. We propose that mutations caused by these substances appear irrespectively of whether cells are dividing or not at the time of exposure. Other agents require their incorporation into the DNA by polymerases as a necessary step to leave a mutational footprint, such as 5-FU/capecitabine and other base analogs^7,43^. Interestingly, their mutations will thus only appear in cells that are not quiescent when the nucleotide pool is altered^29–31,44,45^. This explains why 5-FU mutations do not appear in treatment-related AMLs, and only in some healthy colonic crypts of patients exposed to 5-FU or capecitabine^46^.

One point that remains obscure concerns the potential contribution of platinum-based drugs to the set of driver mutations of treatment-related AMLs. Although mutations in the same genes appear to drive primary and tAMLs, it is still possible that in some cases the mutations driving tAMLs are contributed by the platinum related signature. To solve this issue, more tAML cases from patients exposed to platinum-based drugs need to be sequenced. It is also important to highlight that in this work, we have focused on signatures of single nucleotide variants. It is possible that platinum-based drugs and/or other chemotherapies leave other footprints in the genome in the form of structural variants, some of which might be involved in leukemogenesis.

Exploiting the treatment mutations as a barcode of the clonal expansion of exposed cells, we showed that the original clonal expansion of tAMLs and treatment-related CH differ in their timing with respect to the moment of treatment. While the clonal expansion that founds the tAML is posterior to the moment of treatment, the establishment of the CH clone predates the treatment. In CH, at the time of treatment a variety of small clones of hematopoietic cells already exist. When exposed to platinum-based drugs, HSC clones carrying mutations that hinder the recognition and repair of certain DNA lesions, such as those affecting PPM1D, TP53 or CHEK2 possess an advantage to survive and develop over neighboring clones^15,16,37,38^. Moreover, mutations in different genes are overrepresented across tAML and CH cases. These results suggest that CH and tAML follow different evolutionary pathways. Whether other pre-existing differences at the moment of exposure to the treatment determine one or the other outcome remains to be clarified. The use of treatment mutations as barcodes in the metastatic setting ---that track the strength of the bottleneck--- may be interesting for clinical applications. Clonal neoantigens are more likely to elicit an antitumor immune response^47,48^. Patients with metastases that underwent stronger evolutionary bottlenecks are expected to carry a higher burden of clonal neoantigens ---treatment-related or otherwise--- and thus respond better to immune checkpoint inhibitors.

In summary, our results demonstrate the usefulness of certain mutational signatures, such as those associated with the exposure to chemotherapies to study the evolution of somatic tissues. Furthermore, it opens the door to explore some of the longer-term effects that the exposure to such cytotoxic treatments causes in cancer survivors.

## Methods

### Datasets of tumor somatic mutations

#### A) WGS AML cohort

##### Sequencing in-house treatment-related AML samples

DNA was extracted from six samples of three secondary AML patients. Both samples were bone marrow aspirates corresponding to the diagnosis of the AML and the remission. The short-insert paired-end libraries for the whole genome sequencing were prepared with KAPA HyperPrep kit (Roche Kapa Biosystems) with some modifications. In short, in function of available material 0.1 to 1.0 microgram of genomic DNA was sheared on a Covaris™ LE220-Plus (Covaris). The fragmented DNA was further size-selected for the fragment size of 220-550bp with Agencourt AMPure XP beads (Agencourt, Beckman Coulter). The size selected genomic DNA fragments were end-repaired, adenylated and ligated to Illumina platform compatible adaptors with Unique Dual matched indexes or Unique Dual indexes with unique molecular identifiers (Integrated DNA Technologies). The libraries were quality controlled on an Agilent 2100 Bioanalyzer with the DNA 7500 assay for size and the concentration was estimated using quantitative PCR with the KAPA Library Quantification Kit Illumina® Platforms (Roche Kapa Biosystems). To obtain sufficient amount of libraries for sequencing it was necessary for the low input libraries (0,1 - 0,2 ug) to amplify the ligation product with 5 PCR cycles using 2x KAPA-HiFi HS Ready Mix and 10X KAPA primer mix (Roche Kapa Biosystems). The libraries were sequenced on HiSeq 4000 or NovaSeq 6000 (Illumina) with a paired-end read length of 2×151bp. Image analysis, base calling and quality scoring of the run were processed using the manufacturer#8217;s software Real Time Analysis (HiSeq 4000 RTA 2.7.7 or NovaSeq 6000 RTA 3.3.3).

##### Calling somatic mutations

We obtained 57 whole-genome sequenced AML samples from dbGAP phs000159^24^. Two extra samples of tAML from platinum-treated patients were obtained from EGAD00001005028^25^. The data comprises bone marrow samples at the time of diagnosis and remission. The cram files deposited were reverted to fastqs using bamtofastq^49^. Then, the fastq files from phs000159, EGAD00001005028 and the inhouse cohort were processed in a uniform manner using the sarek^50^ pipeline implemented within nextflow-core (nextflow version 19.10)^51^. Briefly, the pipeline aligns the fastqs to GRCh38 using bwa-mem^52^, and then implements GATK^53^ best practices to mark duplicates and base recalibration, and lastly somatic variant calling. Variant calling of both single nucleotide variants and short insertions and deletions was performed using Strelka2^54^. Only variants labeled as PASS by the algorithm were kept. Variants within regions of low mappability or low complexity^55,56^ were excluded from downstream analyses. All called somatic mutations were annotated with VEP^57^ (version 92).

#### B) WES AML cohort

Within the beatAML cohort^36^, a bone marrow sample and a paired skin sample were taken from 261 AML patients. The somatic mutations identified in these AML cases were downloaded from the Genomics Data Commons^58^ (GDC) repository provided by the authors of the original paper reporting the sequencing and variant calling of these patients. Of note, the original Varscan2^59^variant calling was used, and mutations that coincided with variants present in gnomAD^60^ v2.1 at allele frequency greater than 0.005 were removed. Mutations identified in the 347 remaining patients without a paired normal sample, as well as the clinical information of all patients in the cohort, were downloaded from the cbioportal repository^61^. While the somatic mutations across the 261 cases with paired samples (deemed more reliably true somatic calls) were employed in the driver discovery (see below), mutations identified in AML driver genes across the 608 cases were subsequently taken into account.

#### C) Metastasis cohort

Single base substitutions (SBS) identified across the whole-genome of metastases from 729 Breast, 537 Colon-Rectum, 154 Urinary-Tract and 155 Ovary primary tumors were retrieved from the Hartwig Medical Foundation^32^ (HMF) (DR-110). Somatic mutations (produced by HMF) were processed as explained in our previous publication^5^. Moreover, the WGS of both the tumoral and blood control of the metastatic samples were obtained from the HMF repository for downstream use in the identification of blood somatic mutations. The identification of the somatic mutations in the blood samples of this cohort is explained in Pich *et al*., 2020^38^.

#### D) Healthy blood samples

Whole-genome somatic variants of 23 healthy blood samples were obtained from Osorio *et al*^34^.

### Mutational signatures extraction

Mutational signature extraction in the leukemias in WGS AML cohort, the tumors in the metastasis cohort, and the healthy blood samples in the metastasis cohort was performed using a non-negative matrix factorization approach^62^, as reported in refs^5,38^. We employed the SigProfilerJulia (bitbucket.org/bbglab/sigprofilerjulia) implementation carried out in our lab of the algorithm developed by Alexandrov *et al*^5,63^. The resulting signatures were then compared to the PCAWG COSMIC V3^26^ set using the cosine similarity measure. The Hematopoietic Stem Cell Signature (HSC Sig^34^) was computed as the average the number of mutations observed across the 23 healthy blood samples in each of the 96 tri-nucleotide channels and normalizing them by the total number of mutations observed.

To test for the activity of a mutational signature in a specific sample we used the method so-called mSigAct^39^. Given a set of signatures bound to explain the mutational catalogue of the samples (baseline signatures), the method tests whether an additionally given signature (foreign signature) does improve the sample catalogue reconstruction significantly. Briefly, the method models the mutation count data as being negative binomial distributed and conducts a likelihood ratio test comparing the likelihood of the observed catalogue under two competitive models: with/without the foreign signature. The method returns for each sample the fitting exposures attributed to each signature (both baseline and foreign) alongside the significance (p-value) yielded by the likelihood ratio test.

In the case of the metastatic samples the footprint was deemed as detectable when both SigProfilerJulia and mSigAct report that the treatment signature was active in the treated sample. The set of signatures deemed relevant according to PCAWG^26^ in the respective tumor type cohort was used as a background.

### Driver discovery

The discovery of genes with signals of positive selection in their mutational pattern across the WES AML cohort was carried out using the IntOGen pipeline^64^. Briefly, the IntOGen pipeline integrates seven complementary methods to identify signals of positive selection in the mutational pattern of genes. The IntOGen pipeline first pre-processes the somatic mutations in a cohort of tumors to filter out hypermutators, map all mutations to the GRCh38 assembly of the human genome and retrieve information necessary for the operation of the seven driver detection methods. Then, the methods are executed and their outputs combined using a weighted voting approach in which the weights are adjusted depending on the credibility awarded to each method. Finally, in a post-processing step, spurious genes that result from known artifacts are automatically filtered out. The version of the pipeline used in this study to identify genes under positive selection across the WES AML cohort and the blood samples of the metastasis cohort is described at length at www.intogen.org/faq and in *Martinez-Jimenez et al.*^64^.

To discover drivers across the WES AML cohort, we selected 261 patients with a matched healthy sample, whose somatic mutations were thus considered more reliable. We ran the IntOGen pipeline on the mutations of these samples. Mutations in these or other genes known to drive AMLs^65^ were selected across the 608 patients of the cohort to show in the heatmap of AML drivers

### Clonal and subclonal mutations

We used the MutationTime.R package developed elsewhere^40^ to classify SBS in a tumor as early, late or subclonal. Then, we associated each mutation uniquely with a mutational signature using a maximum likelihood approach^5,66^, and we computed the proportion of clonal mutations amongst all platinum-associated mutations across metastatic tumors.

Subsequently, we computed the fold change between the relative proportions of clonal (grouping early and late clonal mutations) and subclonal mutations associated to the platinum-related signature. We computed this fold change as [(*n*_0_ + *n*_1_) / (*N* 0 + *N* 1)]/ (*n_s_*/*N_s_*), where *n*_0_ and *n*_1_ are, respectively, the number of clonal early and clonal late platinum-related mutations, *n_s_* is the number of subclonal platinum-related mutations, *N*_0_, *N*_1_ and *N_s_* are, respectively, the total number of clonal early clonal late and subclonal mutations.

### Logistic regressions

Multivariate logistic regression analysis was carried out to explore the association between a set of clinical factors and the ability to identify a specific mutational signal associated with chemotherapy. The factors considered were: time of treatment, time since end of the treatment, specific tumor type and whether the metastatic site was considered distal. We considered any metastasis in the same organ as the primary tumor, in a lymph node or in the omentum or peritoneum (in the case of abdominal primary sites) as #8220;proximal#8221; to the primary tumor. Metastases in other sites were considered #8220;distal#8221;. Only tumor types with more than 10 samples were considered for this analysis.

Different multivariate logistic models were considered, each reflecting the sought effect and interactions between the covariates in plausible ways. Model selection was then performed by computing the Bayesian Information Criterion (BIC) and by cross-validation (30% test size, N=100 randomizations) resulting in an average area under the ROC curve (auROC). Model selection gave precedence to models with low BIC and high auROC (Table S2). In both platinum and 5-FU treatments the best model has the same form.

### Compendium of clonal hematopoiesis drivers

A list of 64 genes bearing mutations with signals of positive selection in blood samples was obtained from Pich *et al*., 2020^38^.

### Detection of chemotherapy-associated signatures in blood somatic mutations in the metastasis cohort

We laid out a computational analysis to ascertain the presence of a chemotherapy-induced signature in the blood samples of the metastasis cohort. Given that the low mutation count yielded by the reverse somatic mutation calling precludes a straightforward interpretation sample by sample (high false discovery rate and low statistical power) we proceeded by comparing the distribution of actual reconstructed exposures (inferred from the samples of treated and untreated donors, respectively) against the distribution of exposures reconstructed from synthetic catalogues where the foreign signature has been injected at known levels of exposure.

Given each of 4 possible foreign signatures of interest (associated with platinum and 5-FU)^5^, we generated synthetic mutational catalogues with/without mutations generated by the foreign signature. Next we interrogated the observed and simulated catalogues with a state-of-the-art signature detection method that yields a reconstructed exposure of the foreign signature and a significance level sample by sample^39^. With these outputs we could assess the distributions arising from both observed and synthetic catalogues, thereby assessing the activity of specific mutational processes in a cohort of samples and empirically validating the sensitivity and specificity of the approach. We provide a full description of the methodology in the Supplementary Note.

## Code availability

All analyses described in the paper were implemented in Python and R. All the code needed to reproduce all analyses in the paper will be available via a public repository at the time of publication.

## Data availability

The data employed in the paper is available through different sources. Whole genome sequences of samples in the WGS AML cohort are accessible through the dbGAP phs000159, EGAD00001005028 and EGAXXXXX (to be deposited before publication) collections. Somatic mutations of samples in the WES AML cohort are available through the GDC repository provided by the authors in the original publication of the beat AML cohort (ttps://pubmed.ncbi.nlm.nih.gov/30333627/) and through the cbioportal (www.cbioportal.org). Whole genome sequences of tumor and blood samples in the metastasis cohort are available from the Hartwig Medical Foundation for academic research upon request (https://www.hartwigmedicalfoundation.nl/en).

## Contributions

O.P., A.G.-P. and N.L.-B. designed the project. O.P. carried out all the analyses and prepared the figures. A.C.-B. and M.P. collected the samples and participated in the development of the project. F.M. contributed to the statistical analyses, conducted the analysis of mutational activity with low-count CH data and wrote the Supplementary Note. N.L.-B. and A.G.-P drafted the manuscript. O.P., A.G.-P. and N.L.-B. edited the manuscript. A.G.-P. and N.L.-B. supervised the project.

## Acknowledgements

N.L-B. acknowledges funding from the European Research Council (consolidator grant 682398) and ERDF/Spanish Ministry of Science, Innovation and Universities - Spanish State Research Agency/DamReMap Project (RTI2018-094095-B-I00) and Asociación Española Contra el Cáncer (AECC) (GC16173697BIGA). IRB Barcelona is a recipient of a Severo Ochoa Centre of Excellence Award from the Spanish Ministry of Economy and Competitiveness (MINECO; Government of Spain) and is supported by CERCA (Generalitat de Catalunya). O.P. is the recipient of a BIST PhD fellowship supported by the Secretariat for Universities and Research of the Ministry of Business and Knowledge of the Government of Catalonia, and the Barcelona Institute of Science and Technology (BIST). This publication and the underlying research are partly facilitated by Hartwig Medical Foundation and the Center for Personalized Cancer Treatment (CPCT) which have generated, analyzed and made available data for this research. This work also benefited from DNA whole-exome sequencing data of the BEAT AML cohort, generated within the BEAT AML clinical trial.

---

**Figure S1.**
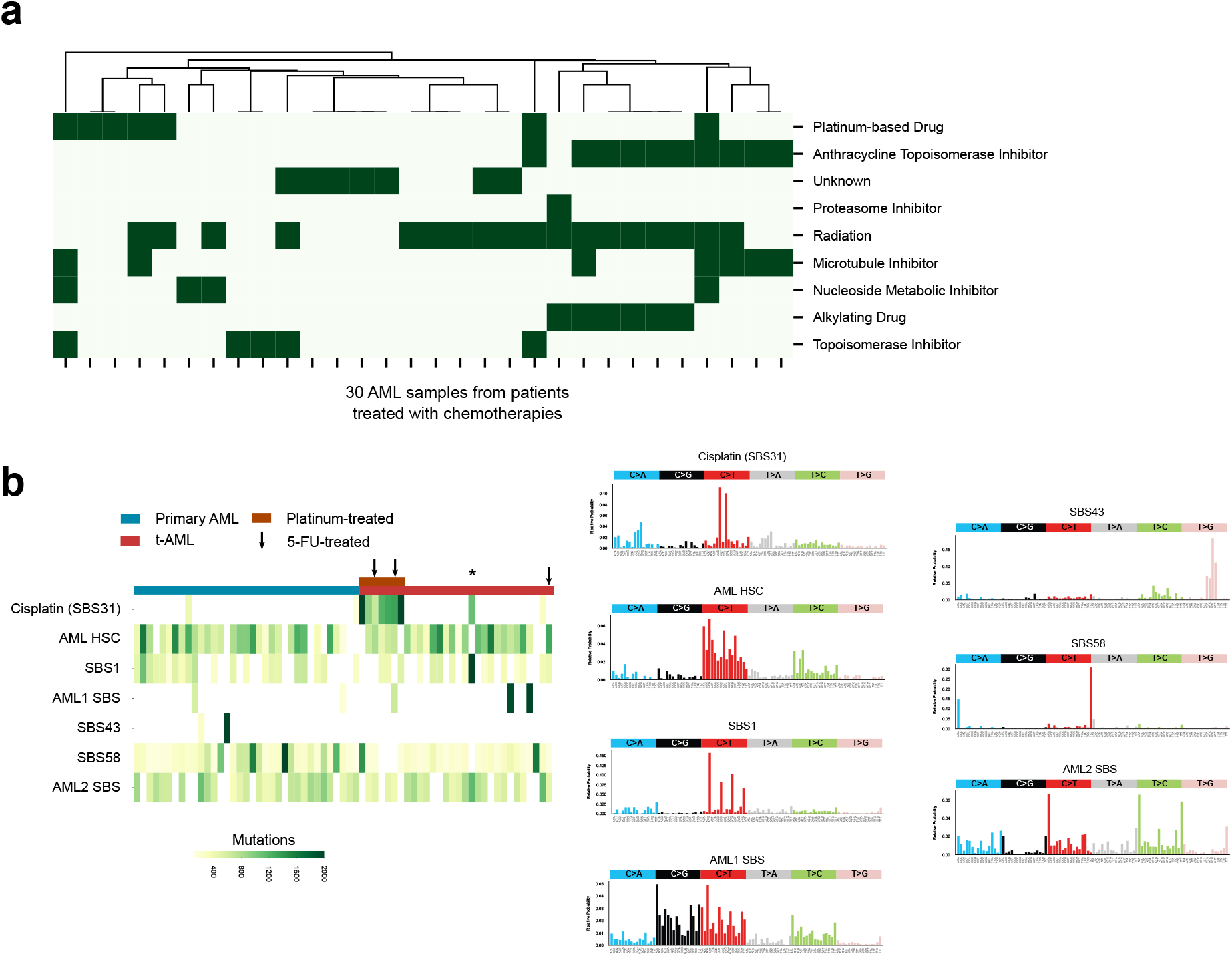
The mutational footprint of chemotherapies across treatment-related AMLs. a)Anticancer treatments to which patients in the WGS AML cohort were exposed as part of the treatment of their primary solid tumors. b)Mutational signatures active across the WGS AML cohort (right panel), and their activity in each sample of the cohort (left). Mutational signatures with known etiology are referred to by their origin, or their number in the compendium of mutational signatures maintained by the COSMIC repository. A t-AML with activity of the platinum-related signature (and a number of double-base substitutions comparable to that of platinum-exposed samples) from a patient who according to their clinical data was not exposed to any platinum-based drug is marked with an asterisk.

**Figure S2.**
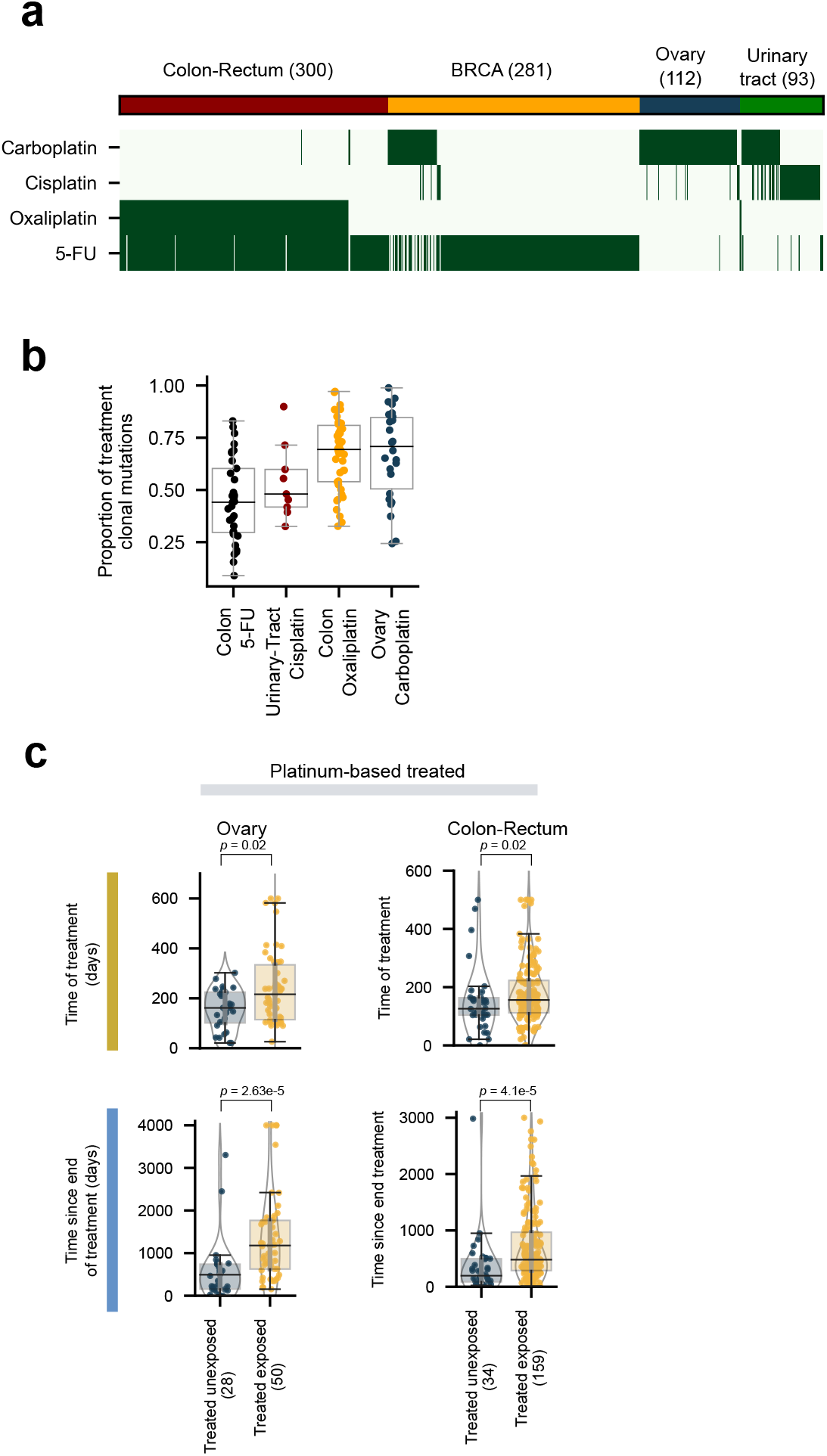
Different mutagenic mechanisms of platinum-based drugs and 5-FU. a)Exposure of metastatic tumors from different organs of origin to platinum-based drugs and 5-FU across the metastasis cohort. b)Distribution of the proportion of clonal treatment-related mutations identified in metastatic tumors from different organs of origin across the metastasis cohort. c)Distribution of time of treatment (days) and time after treatment (days) of samples of metastatic tumors of different organs of origin with detectable or undetectable treatment mutations taken from patients exposed to platinum-based drugs or 5-FU.

**Figure S3.**
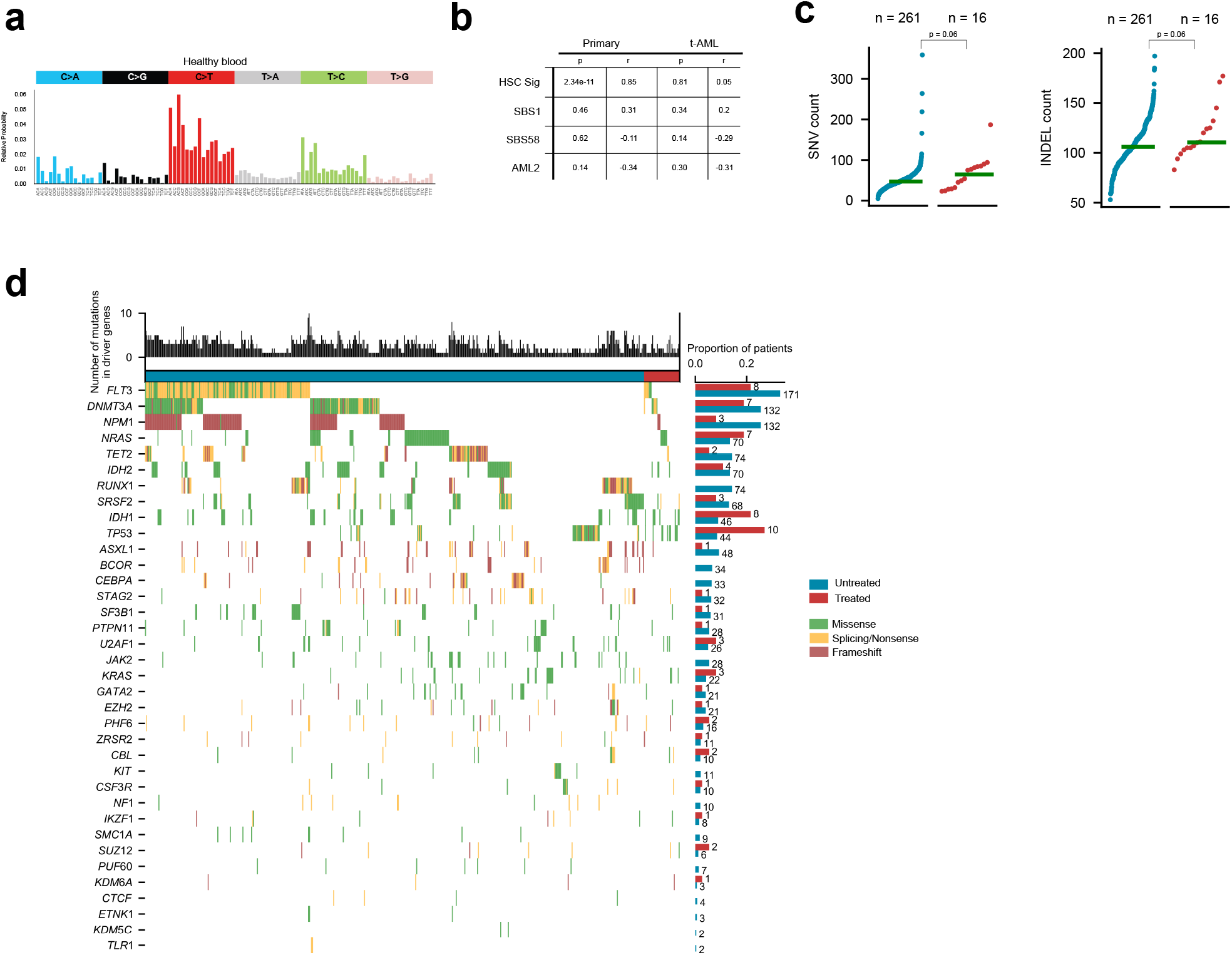
The development of treatment-related AMLs. a)Mutational signature of variants accumulated as a result of hematopoiesis cell divisions, extracted from healthy blood samples^34^. b)Correlation (Pearson#8217;s coefficient and significance) of the number of mutations contributed by all mutational signatures active in primary and tAML cases in the WGS AML cohort and the age of donors. c) Comparison of the number of single nucleotide variants and indels identified in primary and tAML cases of the WES AML cohort. d) Mutations in AML driver genes in primary and tAML cases of the WES AML cohort. The total number of mutations in AML driver genes in each AML patient appears as a bar at the top of the heatmap. The total number of mutations in each gene among primary and tAML cases appears represented as two bars at the right side of the heatmap. Primary and tAML cases are separated in the heatmap.

**Figure S4.**
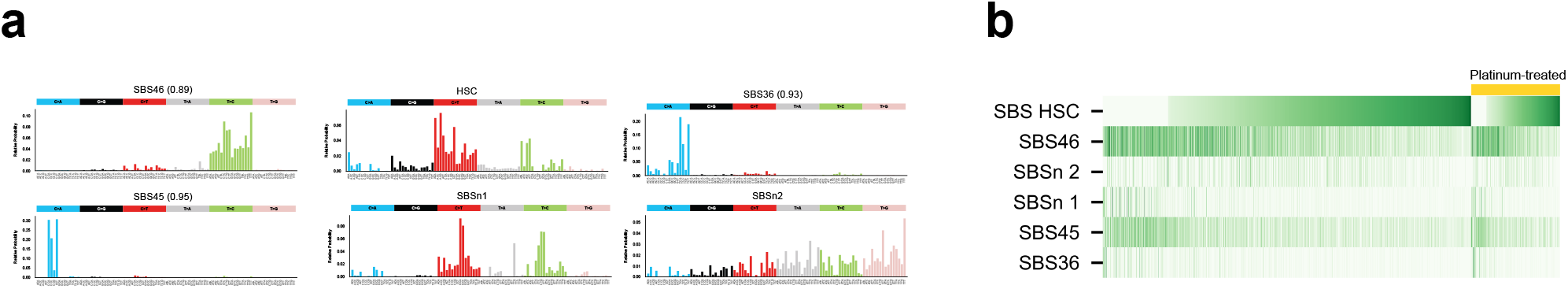
Mutational processes active in normal blood. a)Mutational profile of the six signatures extracted from blood somatic mutations identified across samples in the metastasis dataset. The HSC signature is labeled as such. Signatures related to potential sequencing artifacts are labeled following their number in the COSMIC repertoire of mutational signatures. None of these profiles resembles the platinum-related mutational signature shown in Figure 1d. b)Activity of the six signatures across blood samples of donors in the metastasis cohort. No differences in signature activity are observed across donors exposed and non-exposed to platinum-based drugs.

## Supplementary Tables

**Table S1. Description of the WGS tAML cohort**

**Table S2. Logistic regression model for metastatic tumors exposed to platinum-based drugs and 5-FU**

**Table S3. Treatment-related AML driver genes**

