## Supplementary Note for "The evolution of hematopoietic cells under cancer therapy"

**4 Discussion** **8**

### 1 Summary

When testing for the activity of a mutational process in the catalogue of mutations of a cohort (e.g., signature induced by platinum-based chemotherapy) we can in general resort to signature deconstruction or fitting and statistical testing thereupon, from which we can infer the activity of the signature sample by sample. However, these methods require a high enough mutation count or cohort size to render sufficient evidence.

In our study we were interested in carrying out this kind of signature detection with the requirement to handle low mutation counts: low counts preclude a straightforward interpretation sample by sample because of the high false discovery rate and reduced statistical power.

To circumvent this limitation, we laid out a computational analysis that aims to detect the activity of a foreign signature at the cohort level by applying a signature detection method sample by sample and then pooling the results. The rationale is to compare distributions of reconstructed exposures to reveal latent signature activities in the cohort, even if these activities cannot be confidently distributed across samples.

We generated synthetic catalogues of mutations with/without activity of a specific foreign signature of interest. The approach employed was bound to abstract the common mutational patterns observed in our cohort of study in terms of common mutational processes and exposures.

Next we interrogated the observed and simulated catalogues with a state-of-the-art signature detection method that yields as a result a reconstructed exposure to the foreign signature of interest and a significance level accounting for the reconstruction improvement if the foreign signature is included [1].

With these outputs we compared the observed and simulated catalogues, thereby assessing the activity of specific mutational processes in a cohort of samples and empirically validating the sensitivity and specificity of the signature detection method in our particular setting. We hereto describe the methodological steps in detail and report our findings.

### 2 Synthetic data

We followed closely the simulation approach laid out in [1]. We generated synthetic data sets by the addition of two catalogues of mutations: i) baseline mutations representative of the mutational profile of samples from untreated donors; ii) foreign (chemotherapy-associated) mutations. Here the term *catalogue of mutations* stands for a tabular data set of mutation counts across the 96 possible trinucleotide contexts comprising all possible single base substitutions with a pyrimidine reference allele.

#### 2.1 Baseline-Model Samples

The catalogue of mutations obtained from blood samples of untreated donors of the metastasis cohort (i.e. not subject to any treatment regimen by the time of collection) was taken as the reference for the baseline mutational signal. The starting point is to provide a (96 channel) signature deconstruction of this catalogue. Given the signature deconstruction of the reference catalogue of mutations, we record the proportion $\pi_S$ of samples with non-zero exposure for each active signature $S$ in the deconstruction. Also, for each signature we can compute the best gamma fit for the distribution of non-zero exposures reported for $S$. We denote the resulting gamma distribution $\mathbf{Gamma}(\alpha_S, \beta_S)$. The choice of the gamma distribution choice is justified in order to capture the skewness of the non-zero exposures.

We then use the $\pi_S$ and $\mathbf{Gamma}(\alpha_S, \beta_S)$ computed for each signature $S$ to generate synthetic samples. First, we randomly draw the set of signatures that will be active, independently from each $\mathbf{Bernoulli}(\pi_S)$. Then for each active signature $S$ we draw its exposure $e_S \sim \mathbf{Gamma}(\alpha_S, \beta_S)$. The non-active signatures do not contribute any mutations to the catalogue.

Finally, the mutation count attributed to $S$ in context $c$ is drawn independently from a $\mathbf{NegBinom}(\mu_{S,c}, \sigma)$, the negative binomial distribution with mean $\mu_{S,c}$ and overdispersion parameter $\sigma = 0.1$. The parameter $\mu_{S,c} = e_S \cdot f_{S,c}$ is the mean mutation count generated by signature $S$ in channel $c$, obtained as the product of the exposure of $S$ and the frequency $f_{S,c}$ of context $c$ in the signature $S$. Adding the catalogues generated for all active signatures at each of the 96 contexts, we obtain the catalogue of a synthetic sample.

This procedure to model a typical blood sample from an untreated donor in the metastatic cohort, from which we can randomly draw synthetic catalogues of mutations, has also the practical advantage that it effectively filters spurious signals that may be active in a subset of samples, but are not accounted for by any of the signatures employed in the deconstruction.

#### 2.2 Baseline-Observed Samples

We were also interested in generating synthetic samples with the observed catalogue of mutations of blood samples from untreated donors of the metastasis cohort as baseline.

Therefore, for the subsequent generation of synthetic samples, we resorted to two baseline sets of synthetic samples: so-called baseline-model (4,000 synthetic samples, 1,000 samples independently drawn for each specific foreign signature analysis) and baseline-observed (1,038 blood samples from untreated donors).

#### 2.3 Injecting Foreign Mutations

For this analysis, we probed four previously reported single base substitution signatures associated with platinum-based and 5-FU-based chemotherapy regimens: i) signature E-SBS17b [2] (cosine similarity 0.97 to COSMIC SBS17b), associated to 5-fluorouracil; ii) a signature associated with oxaliplatin; COSMIC (v3.0), iii and iv) signatures SBS31 and SBS35, associated with platinum-based chemotherapy regimens [2, 3]. For this supplementary note, we will label these signatures as: "SBS17b", "oxaliplatin", "SBS31" and "SBS35", respectively.

Synthetic data sets are the result of adding a baseline catalogue and a foreign catalogue, generated from the foreign signature of interest at a specified level of exposure per sample: foreign signatures are thus "injected" in the baseline catalogue. In a similar fashion as described above, if we denote the foreign signature as $S$, injecting $S$ at a level of exposure $e_S$ implies to add as many counts in context $c$ as drawn from $\textbf{NegBinom}(\mu_{S,c}, \sigma)$, the negative binomial distribution with mean $\mu_{S,c} = e \cdot f_{S,c}$ and overdispersion $\sigma = 0.1$.

#### 2.4 Synthetic Catalogues of Mutations

For each signature $S$ out of 4 possible chemotherapy-associated signatures (3 platinum-related and 1 5-FU-related) and for each exposure value in the probing grid

$$\mathcal{G} = \{2, 5, 7, 10, 15, 20, 25, 30, 35, 40, 45, 50, 75, 100\}$$

a synthetic catalogue was generated by taking a baseline catalogue, then injecting the foreign signature to each sample at the exposure value (as described in the previous section). This procedure was carried out with both the baseline-model and the baseline-observed catalogues.

### 3 Foreign Signature Analysis

#### 3.1 Signature Detection Method

To test for the presence of a mutational signature in a specific sample we used the method so-called `mSigAct` [1]. Given a set of signatures bound to explain the mutations of the sample (baseline), the method tests whether the inclusion of an additionally provided signature (foreign) improves the reconstruction significantly.

More specifically, the method models the mutation count data as being negative binomial distributed and carries out a likelihood-ratio test comparing the likelihood of the observed catalogue under two competitive models: with/without the foreign signature. Assuming one of the two competitive models is in place and we have been able to estimate the exposures to each signature that best explain our sample, for each trinucleotide context $c$ we have an actual mutation count $m_c$ and a reconstructed mutation count $r_c$. The log-likelihood of the actual catalogue is then computed as

$$\mathcal{L} = \sum_c \log P(m_c \mid r_c, \sigma),$$

where $P$ is the negative binomial probability of $m_c$, governed by the mean $r_c$ and some convenient overdispersion parameter $\sigma$.

The reconstructed mutation count parameters $r_c$ depend on the estimation of the exposures for the signatures considered in the model (with/without foreign signature). These exposures are obtained upon non-linear optimization, with the likelihood $\mathcal{L}$ acting as objective function. The reader is encouraged to go through the Supplementary Materials and Methods of [1].

Given baseline and foreign signatures, the output of `mSigAct` provides a reconstructed exposure for the foreign signature alongside the significance witnessed by the likelihood-ratio test. Both the synthetic samples obtained from the baseline-model and from the baseline-observed, including the baseline catalogues themselves (with zero mutations injected) were subject to `mSigAct` analysis.

#### 3.2 Observed Samples by Treatment Regimen

We segregated the samples from the metastasis cohort into groups by treatment regimen. For each foreign signature considered we compared the distributions of the most likely reconstructed exposures of the foreign signature upon `mSigAct` analysis and compared the distributions between treatment groups and the baseline group of untreated samples.

We focused on five treatment regimens: carboplatin, cisplatin, oxaliplatin (platin-based); fluorouracil and capecitabine (nucleoside metabolic inhibitors or NMI). For each treatment we conducted treated/untreated comparisons for each foreign signature, thereby interrogating whether the treatment has any discernible effect at the cohort level in terms of the foreign signature activity. For each comparison we carried out a Kolmogorov-Smirnov (KS) two-sample test to render a significance value for the discrepancy between distributions (see Figure SN1 of which the main Figure 6e is the SBS31-cisplatin case). None of the comparisons provided any evidence of significant differences between treated/untreated groups for any of the treatment-associated signatures considered.

We then carried out the same exercise with the distributions of $p$-values upon `mSigAct` analysis. When taking all samples into account, the analysis produced no differences between treated and untreated samples. Then we reasoned that we might still be able to identify a subgroup of samples with a signature-treatment effect if we compared only samples with non-zero reconstructed exposure. We carried out this analysis with different thresholds. In Figure SN2 we show the result for samples with exposure $\geq 5$. None of the comparisons yielded any significant differences between treated/untreated groups for any of the treatment-associated signatures considered. Remark that in one case (signature SBS31 and cisplatin) the comparison showed a weak although non-significant trend favouring the hypothesis that the treated samples bore a footprint of SBS31.

#### 3.3 Synthetic Samples

We also analyzed the synthetic samples with known levels of exposure to injected signatures, including the baseline (zero-injected samples). This analysis allowed us to have a more concrete understanding of what signal-to-noise ratio is expected when considering different levels of exposure to the signature. In other words, considering the previous comparisons, is it possible that at the expected levels of exposure, the treatment-signature association signal was too weak for our statistical procedure to detect it?

For each signature and each level of injection a KS two-sample test was carried out comparing either the exposure or $p$-value distribution of the injected synthetic samples injected to the baseline. We did it in the two settings derived from baseline-observed (Figure SN3) and from baseline-model samples (Figure SN4), respectively.

As expected, in both synthetic data sets the signal-to-noise was dependent on the signature of interest. For example, SBS17b showed the strongest effects, both in terms of reconstructed exposures and $p$-values, even with few injected mutations. On the opposite side, the signal-to-noise induced by SBS31 injection did not reach KS significance ($p < 0.01$) below the level of 7 injected mutations. That said, for all signatures and all levels of exposures, the same trend is apparent.

Interestingly, when comparing the results between both synthetic settings, we can see that the false-positive rate (asserted as the proportion of non-zero exposure reconstructed from zero-injected samples) is lower and more stable in the baseline-model catalogue ($\sim 0.025$) than in the baseline-observed catalogue, where the false-positive rate fluctuates between $\sim 0.125$ (SBS35) and $\sim 0.025$ (SBS17b).

### 4 Discussion

In our analysis we compared treated/untreated samples, then conducted the same signature detection with synthetic samples in two different settings. None of the treated/untreated comparisons rendered significant differences nor trends suggestive of an effect. However with the same tools we were able to identify a clear signal-to-noise in the synthetic data sets, demonstrating that the lack of signal in the first analysis is not due to a technical limitation, but rather to the absence of activity of the foreign signatures.

Only in one case (signature SBS31 and cisplatin) we found a non-significant trend consistent with a small subset of samples having this treatment-signature association.

The analyses and results presented in this supplementary note are highly consistent with the conclusion that there are no homogeneous treatment-signature effects for the signatures and treatments considered across healthy blood samples from patinets of the metastasis cohort.

[2] O. Pich, F. Muiños, M. P. Lolkema, N. Steeghs, A. Gonzalez-Perez, and N. Lopez-

Bigas. The mutational footprints of cancer therapies. *Nat. Genet.*, 51(12):1732–1740, 12 2019.

[3] L. B. Alexandrov, J. Kim, N. J. Haradhvala, M. N. Huang, A. W. Tian Ng, Y. Wu, A. Boot, K. R. Covington, D. A. Gordenin, E. N. Bergstrom, S. M. A. Islam, N. Lopez-Bigas, L. J. Klimczak, J. R. McPherson, S. Morganella, R. Sabarinathan, D. A. Wheeler, V. Mustonen, G. Getz, S. G. Rozen, M. R. Stratton, L. B. Alexandrov, E. N. Bergstrom, A. Boot, P. Boutros, K. Chan, K. R. Covington, A. Fujimoto, G. Getz, D. A. Gordenin, N. J. Haradhvala, M. N. Huang, S. M. A. Islam, M. Kazanov, J. Kim, L. J. Klimczak, N. Lopez-Bigas, M. Lawrence, I. Martincorena, J. R. McPherson, S. Morganella, V. Mustonen, H. Nakagawa, A. W. Tian Ng, P. Polak, S. Prokopec, S. A. Roberts, S. G. Rozen, R. Sabarinathan, N. Saini, T. Shibata, Y. Shiraishi, M. R. Stratton, B. T. Teh, I. V?zquez-Garc?a, D. A. Wheeler, Y. Wu, F. Yousif, and W. Yu. The repertoire of mutational signatures in human cancer. *Nature*, 578(7793):94–101, 02 2020.

**Figure SN1:** Cumulative densities of reconstructed exposures obtained with `mSigAct` from samples of the metastasis cohort. Plots show the cumulative density of the reconstructed exposure distribution of treated versus untreated samples for each treatment (row) and candidate signature (column).

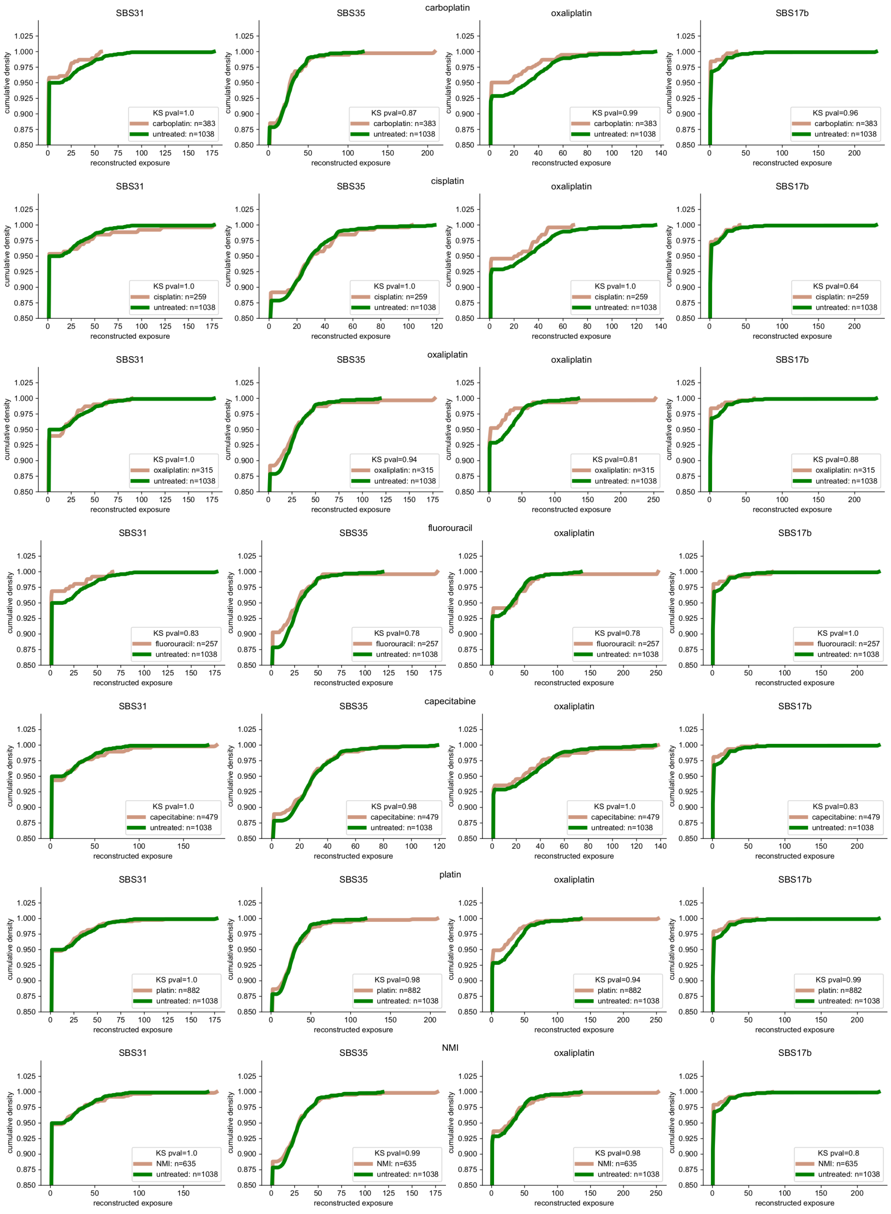

**Figure SN2:** Cumulative densities of reconstructed $p$-values obtained with `mSigAct` from samples with reconstructed exposure $\geq 5$. Plots show the cumulative density of the reconstructed exposure distribution of treated versus untreated samples for each treatment (row) and candidate signature (column).

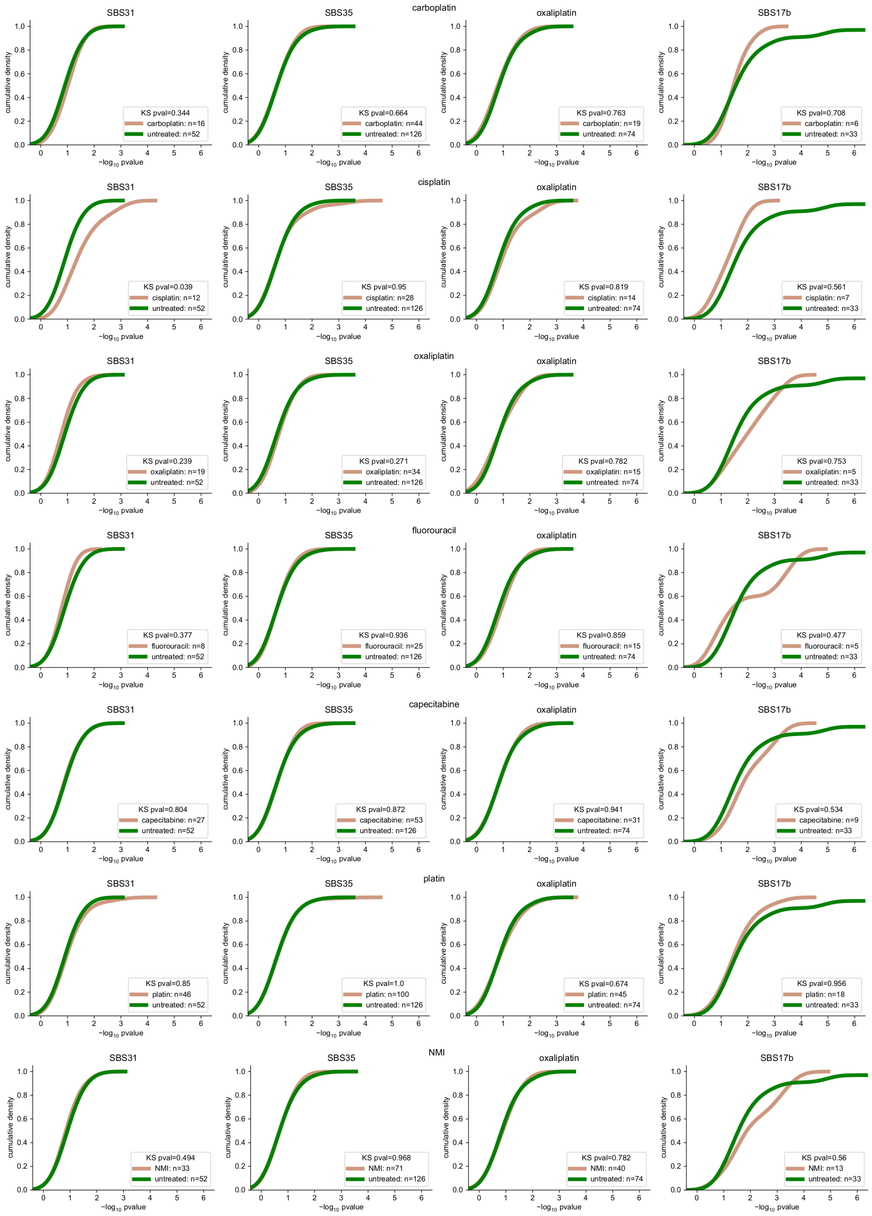

**Figure SN3:** Cumulative density functions of the reconstructed exposures (Panel a) and $p$-values (Panel b) after running `mSigAct` on the synthetic data set derived from the baseline-observed samples.

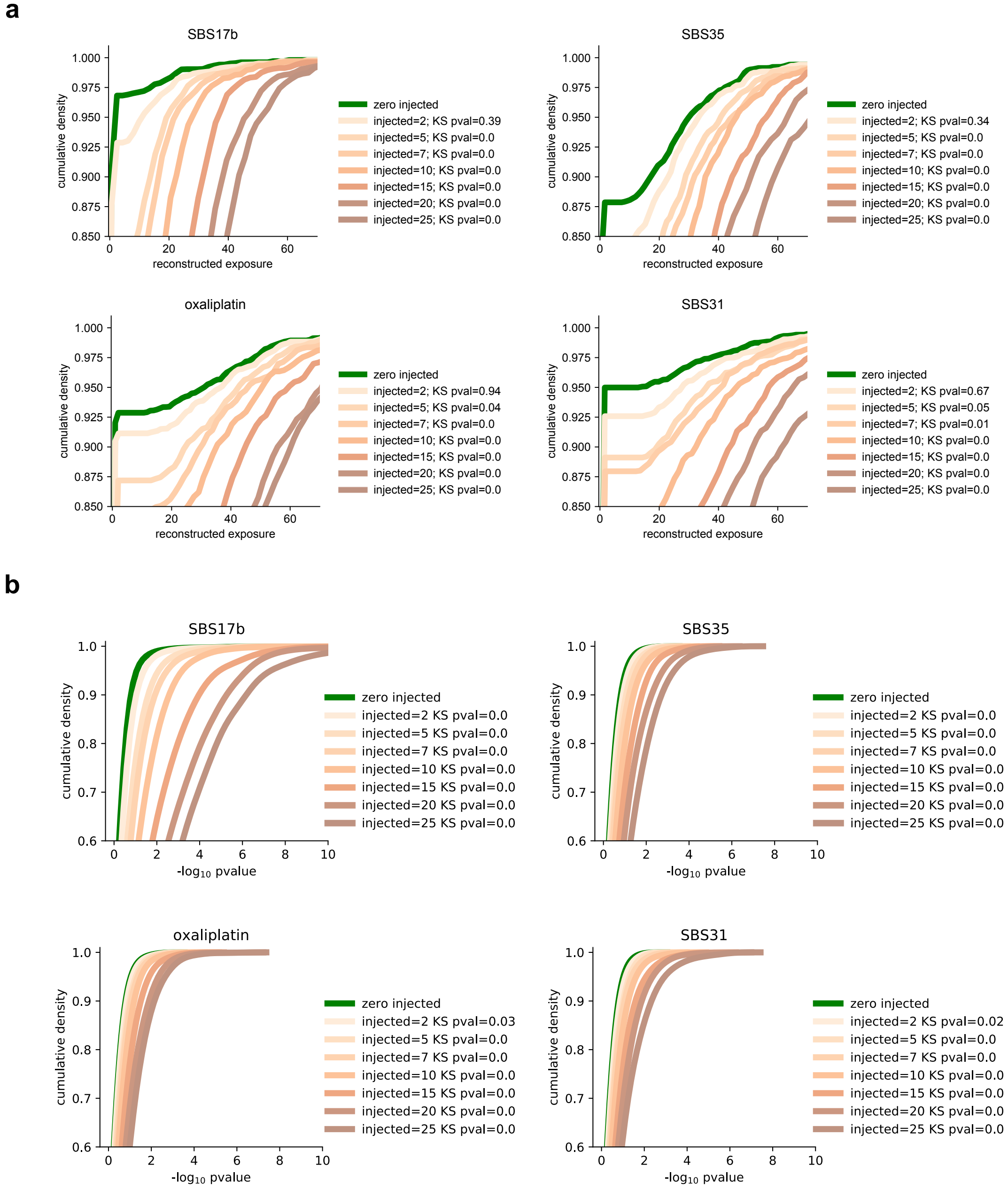

**Figure SN4:** Cumulative density functions of the reconstructed exposures (Panel a) and *p*-values (Panel b) after running `mSigAct` on the synthetic data set derived from the baseline-model samples.

**a**

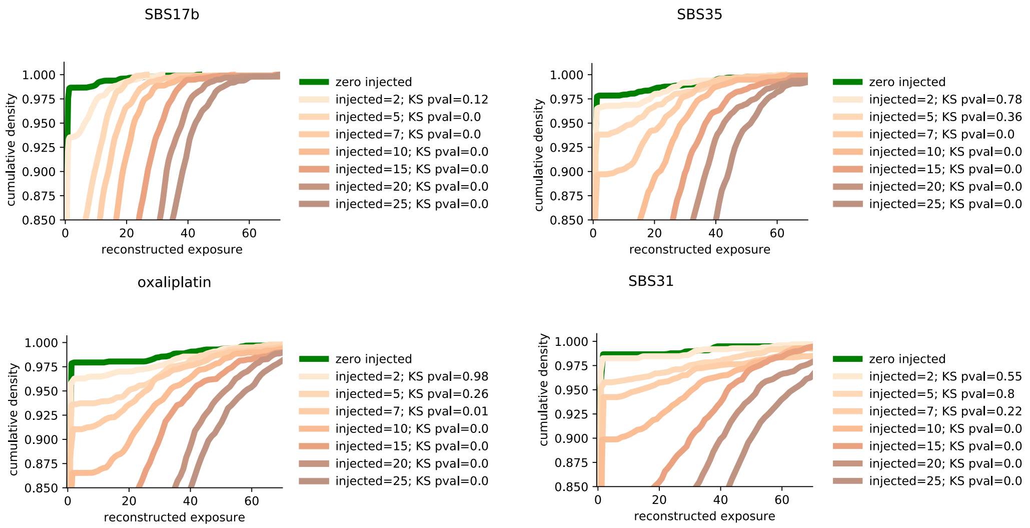

**b**

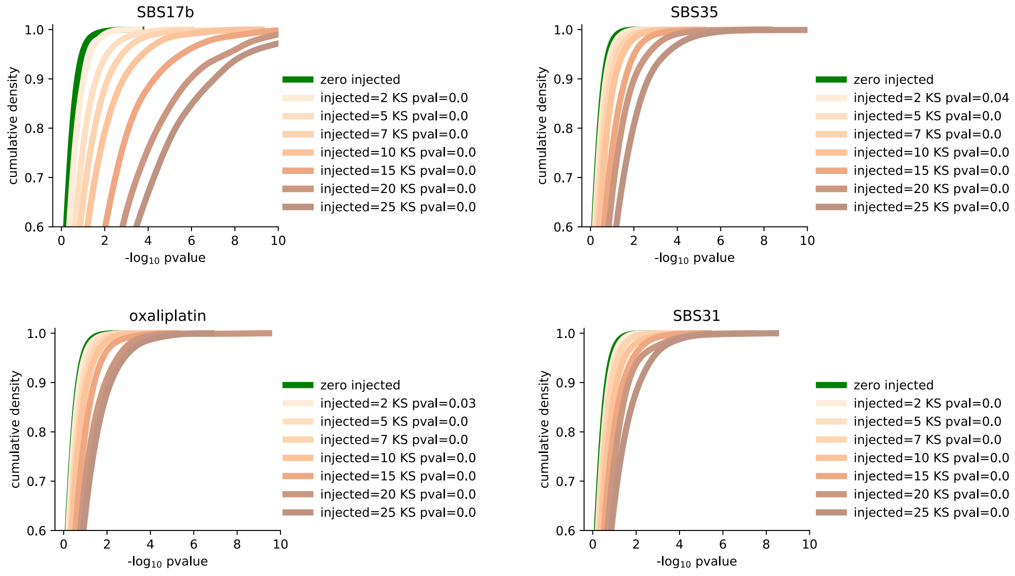
